# Human dorsal root ganglia neuronal cell line to study nociceptive signaling: a new pipeline for pain therapy

**DOI:** 10.1101/2025.09.27.674468

**Authors:** Sara A. Dochnal, Yixing Du, Daniella Bandari, Kaue F. Malange, Jack Bryant, Julia Borges Paes Lemes, Abby Whitford, Anna R. Cliffe, Prashant Mali, Kim Dore, Yury I. Miller, Tony L. Yaksh

## Abstract

Nociceptive afferent neurons within the dorsal root ganglion (DRG) detect and relay painful stimuli from the periphery to the brain, and the malfunctioning of this process leads to sustained pain states. Animal model studies have been invaluable for demonstrating the importance of the DRG nociceptor in pain sensation and the development of related analgesic targets. However, there are functional biological differences between human and animal model nociceptors. Therefore, a complementary *in vitro* model of human nociception is critical to confirming the relevance of preclinical findings for therapeutic drug development. We characterized the nociceptive properties of differentiated cells from the human DRG-derived immortalized cell line HD10.6. Within differentiated HD10.6 cells, we documented the abundance and localization of nociceptive machinery central to regulating excitability and linked with pain sensation including ion channels TRPV1 and NaV1.7 and afferent peptides CGRP and Substance P. Using calcium influx imaging assays, we confirmed the electrical functionality of TRPV1 and NaV1.7 in HD10.6 cells, and through whole-cell patch clamp, we found similar baseline electrophysiological parameters of HD10.6 cells to those previously observed in human patient DRGs. Further, we found that differentiated HD10.6 cells express the mu opioid receptor 1 protein, and DAMGO, a mu agonist, blocks depolarization-evoked calcium influx in a naloxone-reversible fashion. Importantly, using an inflammatory cocktail, excitation and peripheral sensitization are induced within HD10.6 cells, mirroring nociceptors in a pain state during or after tissue damage or inflammation. Finally, HD10.6 cells were also cultured into dual-chambered microfluidic devices to mirror the biological anatomy of the nociceptor. Within this system, we demonstrated the uptake of adeno-associated-virus (AAV) by the peripheral terminals and AAV transport to the soma. Altogether, we have developed the use of HD10.6 cells to create a system of human nociceptive signaling on a chip to study human nociceptor physiology and intervention.

**Perspective:** There are essential differences between human and animal model nociceptors. Here, we develop a physiological model of “nociceptive signaling on a chip” using human-derived nociceptors to ultimately enhance the translatability of preclinical afferent signaling research to the human patient.

## Introduction

Following peripheral tissue injury and inflammation, afferent signaling has been shown to underlie many aspects of the consequential pain phenotype. Decoding the physiology and disruption of afferent signaling is essential to the creation of therapeutics that selectively regulate the pain state they subserve. Thus far, rodents have represented invaluable tools for studying afferent signaling. *In vivo* rodent models offer behavioral pain readouts and cultured primary rodent DRG neurons, some of which are immortalized into cell lines^1^, have long offered a reductionist *in vitro* approach to studying the molecular workings of nociceptors. While many physiological nociceptive pathways are conserved between rodents and humans^2,3^, there are nonetheless essential anatomical, cellular, molecular, transcriptional, and electrophysiological differences between DRG neurons from animal models and humans (for review, we recommend^3,4^). Species differences, particularly those transcriptomic in nature, have been recently highlighted by a series of publications that have taken high resolution sequencing approaches to human DRGs^5–9^. The expression patterns of certain genes implicated in pain signaling are not equal between species; for example, a comparative genomic study revealed that mu opioid receptor (MOR) expression enriched in peptidergic nociceptors is much greater in humans than rodents. Other genes expressed in peptidergic nociceptors are specific to humans and lack any equivalent in rodents or non-human primates. These differences may have profound implications for translating data from rodent models to human pathologies, and subsequent therapeutic developments. In view of the large-scale failure of clinical trials based on animal models^10^, the success of new drugs to treat pain in the clinic will likely require complementary studies of human cells and tissues.

In 1999, Raymon *et al*. created a human-derived immortalized DRG neuronal cell line entitled “HD10.6”^11^. HD10.6 cells are derived from DRGs and immortalized with a tetracycline-inducible c-myc retroviral vector, resulting in a passable clone of neural precursors. Following the addition of tetracycline to suppress c-myc expression and several growth factors to facilitate differentiation, precursors develop into cells with sensory neuron properties. Differentiated HD10.6 cells transmit action potentials^11^ and demonstrate immunoreactivity for several filaments, receptors, and transcription factors characteristic of DRG sensory neurons^11–13^. Therefore, HD10.6 cells have the profound potential to provide species-specific insight into DRG neuron physiology. However, the use of this cell line has been limited since its introduction. HD10.6 cells require further characterization at the RNA, protein, and functional level to evaluate their suitability as a nociceptive system for studying peripheral pain sensation and further optimization to maximize their utility to ultimately to be used as a drug pipeline assay.

Here, we begin by characterizing HD10.6 cells as *de facto* human nociceptors. We highlight the abundance and behavior of characteristic nociceptive ion channels, excitatory/inhibitory receptors, and afferent peptides, including Transient Receptor Potential Vanilloid 1 (TRPV1), Sodium channel voltage-gated NaV1.7, calcitonin gene-related peptide (CGRP), Substance P, neurokinin-1 receptor (NK1R), and MOR. Using whole-cell patch clamp, we reveal similar electrophysiological parameters like those observed in human patient DRGs. Importantly, we induce excitability in HD10.6 cells using an inflammatory cocktail to mirror nociceptors in a pain state during or after tissue damage or inflammation. HD10.6 cells were also cultured into dual-chambered microfluidic devices, in which we can isolate the soma and axon terminals of the nociceptor and mirror the nociceptor’s biological anatomy. HD10.6 cells display successful AAV infection following administration to the peripheral terminal, mirroring the peripheral delivery of pain therapeutics. Altogether, we have used HD10.6 cells to effectively model nociceptor physiology and therapeutic intervention.

## Materials & Methods

### Reagents

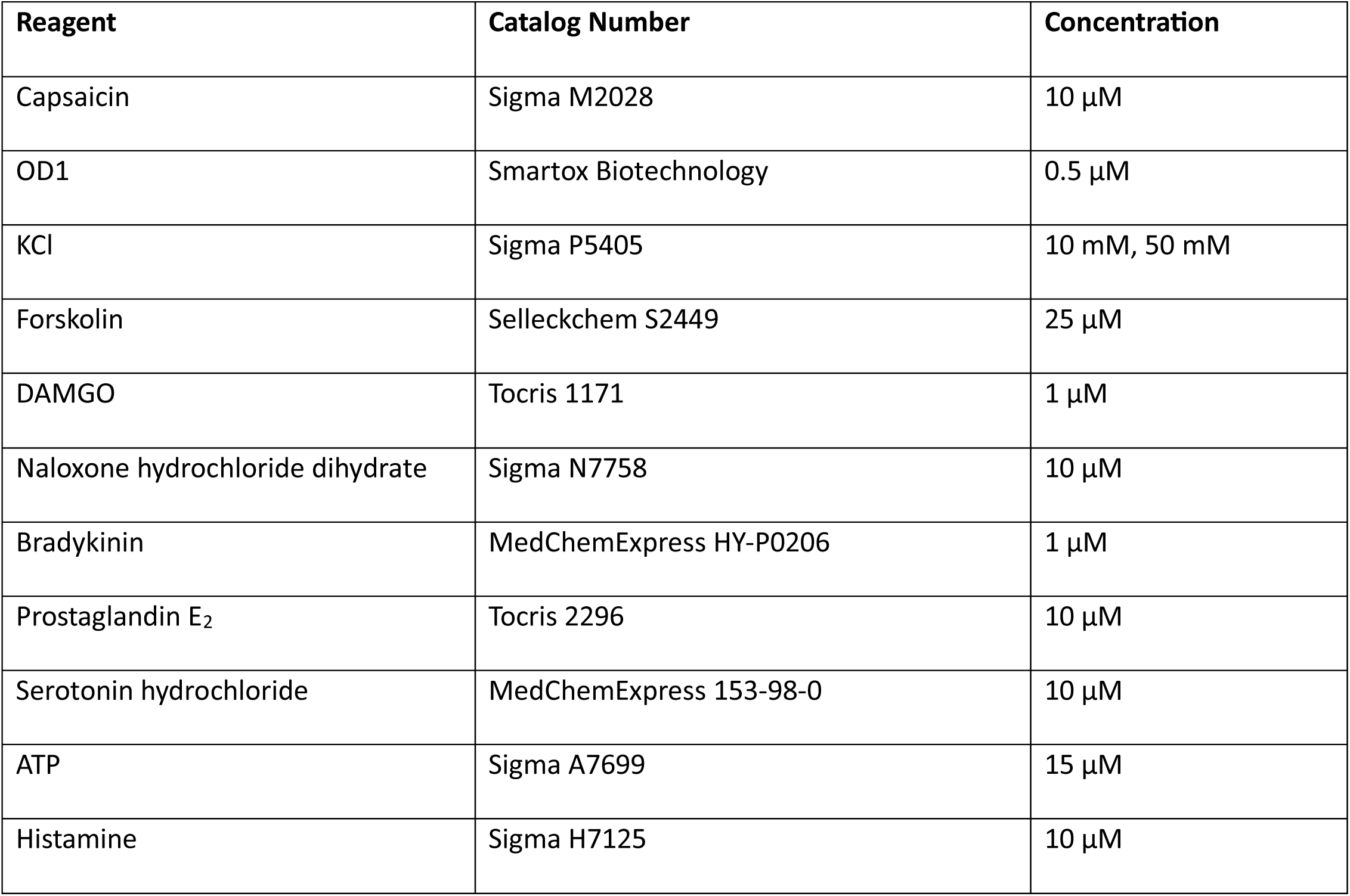

| Reagent | Catalog Number | Concentration |
| --- | --- | --- |
| Capsaicin | Sigma M2028 | 10 μM |
| OD1 | Smartox Biotechnology | 0.5 μM |
| KCl | Sigma P5405 | 10 mM, 50 mM |
| Forskolin | Selleckchem S2449 | 25 μM |
| DAMGO | Tocris 1171 | 1 μM |
| Naloxone hydrochloride dihydrate | Sigma N7758 | 10 μM |
| Bradykinin | MedChemExpress HY-P0206 | 1 μM |
| Prostaglandin E<sub>2</sub> | Tocris 2296 | 10 μM |
| Serotonin hydrochloride | MedChemExpress 153-98-0 | 10 μM |
| ATP | Sigma A7699 | 15 μM |
| Histamine | Sigma H7125 | 10 μM |

### HD10.6 Cell Culture

All cells were incubated at 37°C and 5% CO_2_. Proliferation and differentiation protocols were guided by previous publications^11,12^. Immortalized human dorsal root sensory ganglion cells (HD10.6 cells^11^) were passaged in proliferation media on Nunc flasks (Thermo Fisher Scientific 153732) coated for 3 hours with 17 μg/μL fibronectin (Thermo Fisher Scientific 33016-015) in 1X PBS. Proliferation media consisted of Advanced Dulbecco’s modified Eagle medium (Gibco Life Technology 12634010) supplemented with glutaMAX (Gibco 35050061), Prime-XV IS21 (Fujifilm 91142), and 10 ng/mL prostaglandin E1 (Sigma P5515). Fresh 0.5 ng/mL fibroblast growth factor-basic (bFGF) (Stemgent 03-0002) was added prior to media change.

For differentiated HD10.6 cells, glass coverslips (Fisher Scientific) were prepared in advance to enhance adherence. Coverslips were coated with 10 μg/mL Poly L-ornithine (Sigma P4957) in borate buffer (Boston Bioproducts BB-66) overnight at 37°C and washed three times with sterile water (Millipore Sigma W3500) the following morning. The coverslips were then coated with 1 μg/mL fibronectin in sterile water overnight at 37°C, washed three times with sterile water the next morning, and left to dry for within the hood for 24-72 hours. Cells were seeded on day 0 (D0) at a density of ∼16,000 cells/cm^2^ in proliferation media on coated coverslips. Media was changed to complete differentiation media 24 hours after plating (D1), with half-volume medium changes performed every 2 days afterwards. Complete differentiation media consisted of Neurobasal - phenol red -L-Glutamine (Gibco-Life Tech C12348-017) supplemented with glutaMAX (Gibco 35050061), Prime-XV IS21 (Fujifilm 91142), 50 ng/mL 2.5S nerve growth factor (NGF) (Alomone labs N-100), 25 ng/mL of ciliary neurotrophic factor (CNTF) (PeproTech 450-13), 25 ng/mL glial cell-derived neurotrophic factor (GDNF) (Alomone Labs G-240), and 25 ng/mL neurotrophin-3 (NT-3) (PeproTech 450-03). Fresh 1 μg/mL doxycycline (Sigma T-7660) was added prior to media change.

### Immunofluorescence

HD10.6 cells on coated 12 mm circle glass coverslips were washed twice with 1X PBS and fixed for 10 minutes on ice with 4% paraformaldehyde (PFA). Cells were blocked using 2% fetal bovine serum (FBS), 0.2% Triton-X, and 2% serum from the appropriate host species (donkey or goat) of secondaries for 1 hour at room temperature. Cells were incubated with primary antibody diluted in blocking buffer for 2 hours at room temperature. Following three 1X PBS washes, coverslips were incubated with secondary antibody diluted in blocking buffer for 1 hour at room temperature, covered from light. Following three 1X PBS washes, coverslips with cells were mounted on slides with ProlongGold containing DAPI (Thermo Fisher) and imaged using a Leica SP8 microscope. Images were analyzed using ImageJ.

| Antibody | Catalog Number | Dilution |
| --- | --- | --- |
| NF200 | Millipore AB5539 | 1:1000 |
| β3-Tubulin | Millipore AB9354 | 1:100 |
| TRPV1 | Novus Biologicals NBP1-97417 | 1:200 |
| NaV1.7 | Alomone Labs ASC-008 | 1:100 |
| CGRP | Sigma Aldrich C8198 | 1:100 |
| Substance P | Merck Millipore A1566 | 1:80 |
| NK1R | Merck Millipore AB5060 | 1:80 |
| MOR1 | Abcam ab10275 | 1:80 |
| ATTO488-ProTx-II | Smartox | 1:200 |
| mCherry | Abcam ab167453 | 1:500 |

### Calcium Imaging

All calcium imaging experiments were performed on differentiated HD10.6 cells plated on coated 15 mm glass coverslips. HD10.6 cell cultures were loaded with the Ca^2+^ indicator Fluo-4 AM (5 μM; Invitrogen F23917) with PowerLoad Concentrate (Thermo Fisher P10020) in differentiation media without doxycycline for 1 hour at room temperature. Coverslips were placed in a laminar flow perfusion chamber (Warner Instrument Corp., UK) and continuously perfused with extracellular solution (in mM: NaCl 160; KCl 2.5; CaCl2 1; MgCl2 2; HEPES 10; glucose 10; pH 7.4). The perfusion system consisted of up to four individual compartments, filled with treatment solutions, each one connected to a separate cannula that merged to one output cannula. The latter was inserted in the laminar flow perfusion chamber containing a glass coverslip with the HD10.6 cells. The flow was constant and the HD10.6 cells were always covered in solution. Each treatment valve was opened manually by the experimenter. A suction pump connected to outflow side of the chamber maintained a continuous perfusion. HD10.6 cells were recorded in an inverted Leica TCS SP5 confocal microscope. One field of view per coverslip was assessed. Cells were exposed to buffer, then stimulated with drug diluted in buffer with times of exposure as specified within figure legends. All cultures were stimulated with 50 mM KCl for 5 seconds at the end of experiments to confirm cellular viability. Neuronal Ca^2+^ responses were analyzed in 3–5 dishes per biological replicate by selecting individual cells as ROIs (Region of Interest) and calculating mean gray value variations on each individual cell using LAS AF version 2.7.3.9723 software. Data are presented as ΔF/F0, where F0 is the baseline, and the effect is quantified as maximal ΔF/F0.

### Electrophysiological recordings

Whole-cell patch clamp was performed on intact, differentiated HD10.6 cells plated on coated 15 mm circle glass coverslips. Coverslips were transferred into the recording chamber with a continuous flow of extracellular solution containing 150 mM NaCl, 4.0 mM KCl, 2.0 mM CaCl_2_, 2.0 mM MgCl_2_, 10 mM glucose, and 10 mM HEPES (pH 7.4). The extracellular solution was perfused at a rate of 1.5-2.0 mL/minute and heated to 30-32 °C with a temperature controller (TC-324B, Warner Instrument). The cells were allowed to rest in the chamber for 5-10 minutes before all recordings. HD10.6 cells were identified under the microscope of the recording rig by their small, oval shape cell body (**Fig. 3A**). Borosilicate glass pipettes (outer diameter: 1.5 mm, Warner, Hamden, CT) were pulled from a micropipette puller (Model P-97, Sutter). The recording electrodes had a resistance of 2-5 MΩ when filled with an internal solution containing 130 mM potassium gluconate, 1.0 mM MgCl_2_, 1.0 mM CaCl_2_, 10 mM HEPES, 4 mM MgATP, 0.3 mM Na_2_GTP, and 5.0 mM EGTA at pH 7.2. A MultiClamp 700B amplifier, an Axon Digidata 1550B, and Clampex 11 software (Molecular Devices, San Jose, CA, USA) were used for data acquisition, digitized at 2-10 kHz, and filtered at 2 kHz. The liquid junction potential was compensated for prior to forming the cell-attached mode for all recordings. A minimum of 2 GΩ seal resistance was required before rupturing the membrane for whole-cell configuration. Membrane test in Clampex software was used to read the membrane capacitance (Cm), membrane resistance (Rm), and access resistance (Ra) immediately after whole-cell configuration. The cells were recorded in current clamp mode first to acquire resting membrane potential (resting Vm) without holding current, rheobase (10 ms duration of minimal current injection from 0 pA with 10 pA increment until the first action potential appears), and excitability (500 ms duration of -50 to +500 pA current injection steps with 50 pA increment). Then the cells were held at -70 mV in voltage clamp. When the holding current stabilized, a voltage step protocol was applied to record the Na^+^ currents and K^+^ currents. Voltage steps were from -200 to +40 mV with 10 mV increment and 25 ms duration, and with the baseline holding at -80 mV. Only the cells with Ra <= Rm x 10% and less than 20% fluctuation of Ra were included for voltage clamp recordings. Multiple cells were recorded from each coverslip; and each coverslip were kept in the recording chamber for less than 2 hours to ensure that the cells were recorded in a healthy state.

All the data were analyzed in software ClampFit (Molecular Devices). For excitability analysis, “Threshold search” function ClampFit was used, the depolarization peaks with overshoot above 0 mV were identified as action potentials, and only action potentials occurring during the current injection period were included for quantification of induced action potentials. For the cells showing spontaneous firing at rest, action potentials occurring outside the range of current injection period were not included for induced action potential analysis. For Na^+^ currents analysis, leak current subtraction was performed in ClampFit to isolate depolarization steps induced inward Na^+^ currents, the peak amplitude of each step was used to plot I-V curve, and the maximum of the peak amplitude was used to calculate the current density. The outward currents induced by depolarizing steps near the end of voltage steps were used for K^+^ current I-V plot, and the currents at the +40 mV step were used to calculate the K^+^ current density.

### Microfluidic Chambers

Methods of chamber preparation and use were based off previous publications^14,15^. 24 x 30 rectangle glass coverslips were first exposed to a flame following submersion in 90% ethanol. They were then soaked in 70% ethanol for an hour with constant agitation, dried in the laminar flow hood, and then exposed to UV light for 30 minutes on each side. Sterilized coverslips were coated for differentiation as above with PLO and fibronectin. Silicone microfluidic devices with a 450 μm microgroove barrier (Xona; SND450) were soaked in 70% ethanol for an hour with constant agitation, dried in the laminar flow hood, and then exposed to UV light for 30 minutes on each side. Sterilized microfluidic devices were then adhered to coated coverslips with gentle pressure. On D0, 40,000 HD10.6 cells were seeded into the soma compartment in 20 μL proliferation media and incubated at 37°C and 5% CO_2_ for 10 minutes, prior to topping off media in the soma compartment. The following day, media within the soma compartment was changed to differentiation media and differentiation media was added with a 2X NGF concentration to the axon-only compartment. Half media changes were performed every 2 days for each compartment.

### AAV9-mCherry Production

AAV was produced in house by triple-transfection of HEK293T cells and purified with an iodixanol gradient as previously described^16^. 56 hours prior to transfection, cells were seeded at 10% confluency to achieve 80% to 90% confluency at transfection. Media was exchanged completely 2 hours pre-transfection and 16 hours post-transfection. 15 cm plates were transfected with 30 μg total of transgene vector (mCherry), capsid vector (pXR-9), and pHelper vector in an equimass ratio utilizing linear polyethylenimine (PEI) at a ratio of 4:1 PEI to DNA (PEI dissolved 1 mg/mL in DPBS, pH balanced to 7 with HCl and NaOH). OptiMEM (ThermoFisher) was used to promote complex formation at a final volume of 500 μL transfection mixture per plate. Mixture was briefly vortexed then incubated at room temperature for 10 minutes before dropwise addition to cells. 84 hours after transfection, virus was harvested from supernatant via overnight 4°C 10% polyethylene glycol (PEG) incubation and directly from freeze-thaw lysed cells. Benzonase (SigmaAldrich) incubation at 37°C for 1 hour was used to digest any unencapsulated DNA. An iodixanol gradient was used to isolate virus which was then dialyzed using 50 kDa MWCO centrifugal filters (Millipore) in a solution of DPBS (ThermoFisher), 50 mM NaCl, and 0.0001% of Pluronic F68 (ThermoFisher) and concentrated to a final volume of ∼200 μL. Viral titers were quantified via qPCR of serial dilutions using a known standard (ATCC VR-1616) and primers targeting the ITR regions: AAV-ITR-F (5’-CGGCCTCAGTGAGCGA-3’) and AAV-ITR-R (5’-GGAACCCCTAGTGATGGAGTT-3’).

### AAV9-mCherry Transduction

Within individual wells of a 24-well plate, 1E8, 1E9, or 1E10 vg (viral genomes) of AAV9-mCherry were added in 200 μL of maturation media without doxycycline and incubated for 2 hours at 37°C and 5% CO_2_, following which 400 μL of media with doxycycline was added to reservoirs. Within microfluidic chambers, 1E9 vg were added in 5 μL of maturation media without doxycycline into the axon-only compartment and incubated for 2 hours at 37°C and 5% CO_2_, followed by the addition of media with doxycycline to reservoirs. A 100 μL volume differential was always maintained between axon and soma compartments.

### Statistical Analyses

Results were analyzed using GraphPad Prism v.10 software (GraphPad®, San Diego, USA). The normal distribution of values in each analysis was performed with the Kolmorogov-Smirnov test according to the sample size. When a non-parametric statistical test was performed, the Mann-Whitney test was used to compare two means. When the comparison involved more than two means, the Kruskal-Wallis test was performed to determine if there were differences between the groups analyzed. If the values followed a normal distribution, parametric statistical tests were performed. The T-student test was selected when two means were compared. When the comparison involved more than two means, a one-way or two-way analysis of variance (ANOVA) was performed according to the experimental design followed. The level of significance adopted in each analysis of the results was p < 0.05. Specific analyses are included in the figure legends.

## Results

### Key ion channel expression, localization, and function

To confirm the neuronal origin and nature of HD10.6 cells, we stained proliferating, undifferentiated HD10.6 cells (day 0, D0) and differentiated (10 days-post differentiation, D10) HD10.6 cells for canonical markers of neurons (**Figure 1**). Sensory^17,18^ neuron neurofilament NF200 was detected abundantly and specifically within the cytoplasm of all D0/D10 HD10.6 cells (100%, **Fig. 1A**) as identified by the presence of DAPI-positive staining. Therefore, we proceeded to use NF200 as a positive control for staining in D0 and D10 cells. The presence of neuronal markers in D0 cells despite their relative immaturity is not surprising; immunoreactivity for neuronal microtubule component β3-tubulin^11,12^ has previously been demonstrated in both D0 and D10 cells and likely reflects the neuronal lineage-commitment of the HD10.6 cell line. We also confirmed the presence of pan-neuronal markers β3-tubulin and PGP9.5 within D0/D10 HD10.6 cells (data not shown). The structural nature of NF200 highlights the morphology of undifferentiated and differentiated HD10.6 cells. While D0 HD10.6 cells exhibit a fibroblast-like morphology, rounded soma and axonal projections were readily visible by D4 post-differentiation and became more distinct over time.

**Figure 1:**
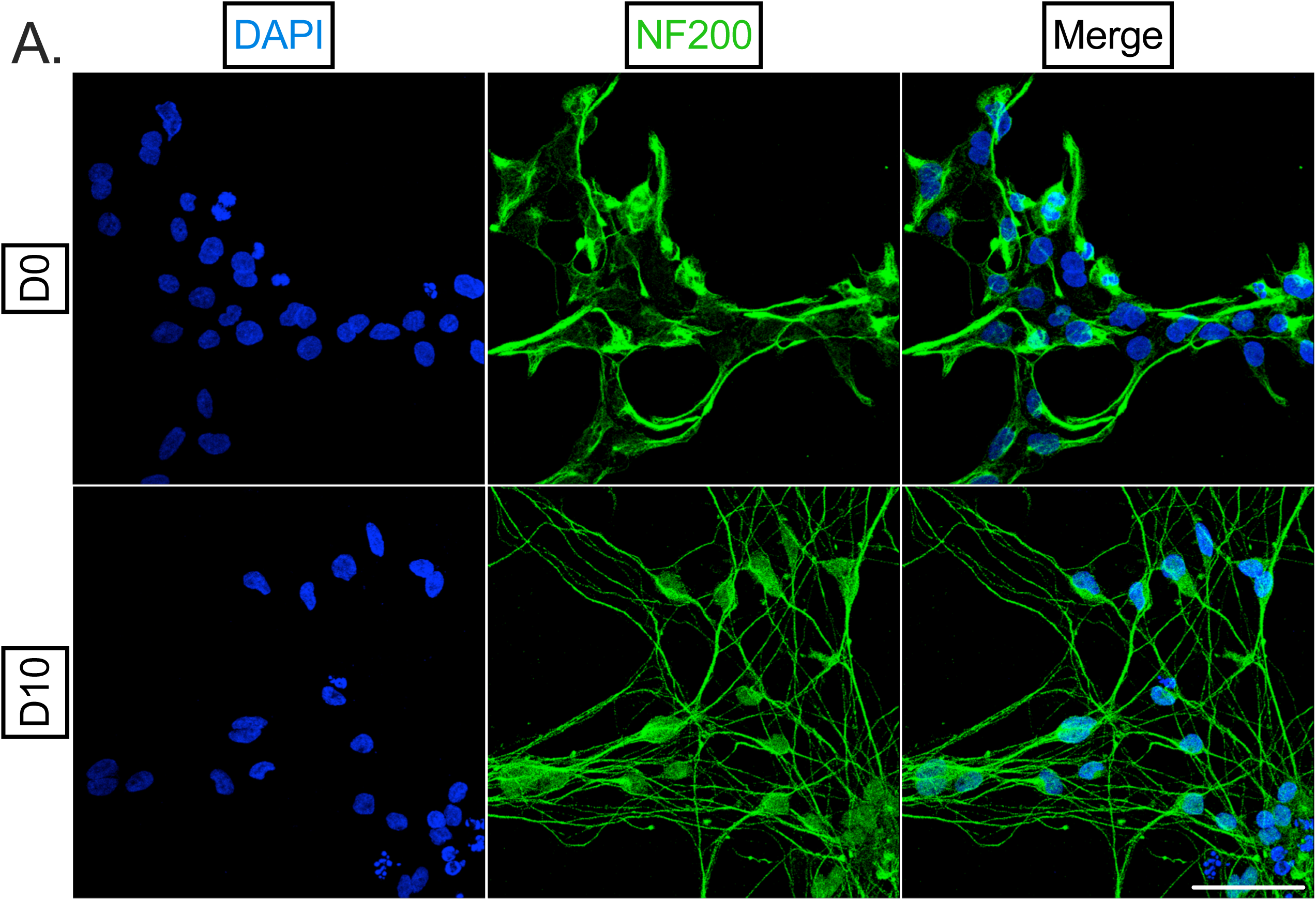
HD10.6 neuronal morphology. **A**) Undifferentiated (D0) and differentiated (10 days post-differentiation, D10) HD10.6 cells were fixed and stained for nuclear stain DAPI (blue) and neurofilament NF200 (green). Scale bar 50 μm. Representative images shown from three independent experiments.

The electrical excitability of nociceptors is regulated by ion channels, the activation of which enables neuronal depolarization, action potentials, and communication of stimuli. Select ion channels within nociceptors have been implicated in the initiation or persistence of pain states, and therefore their expression and function in this system is essential to investigate (**Figure 2**). Paradigm ion channel, non-selective cation channel TRPV1, plays a crucial role in pain and temperature sensation and can be activated by an array of stimuli, including capsaicin, heat, and low pH. Within D0 and D10 HD10.6 cells, we detected immunoreactivity for TRPV1, with localization of TRPV1 to both the soma (filled arrows) and axons (empty arrows) of differentiated HD10.6 cells (**Fig. 2A**). No signal was detected in control slides incubated without primary antibodies. To confirm the electrical functionality of TRPV1 within the HD10.6 cells as previously reported^11^, we tracked calcium influx events by live imaging following exposure to capsaicin. Acute (30s) exposure of HD10.6 cells to 10 μM capsaicin induced a transient and statistically significant increase in influx representative of an action potential (**Fig. 2C-D**).

**Figure 2:**
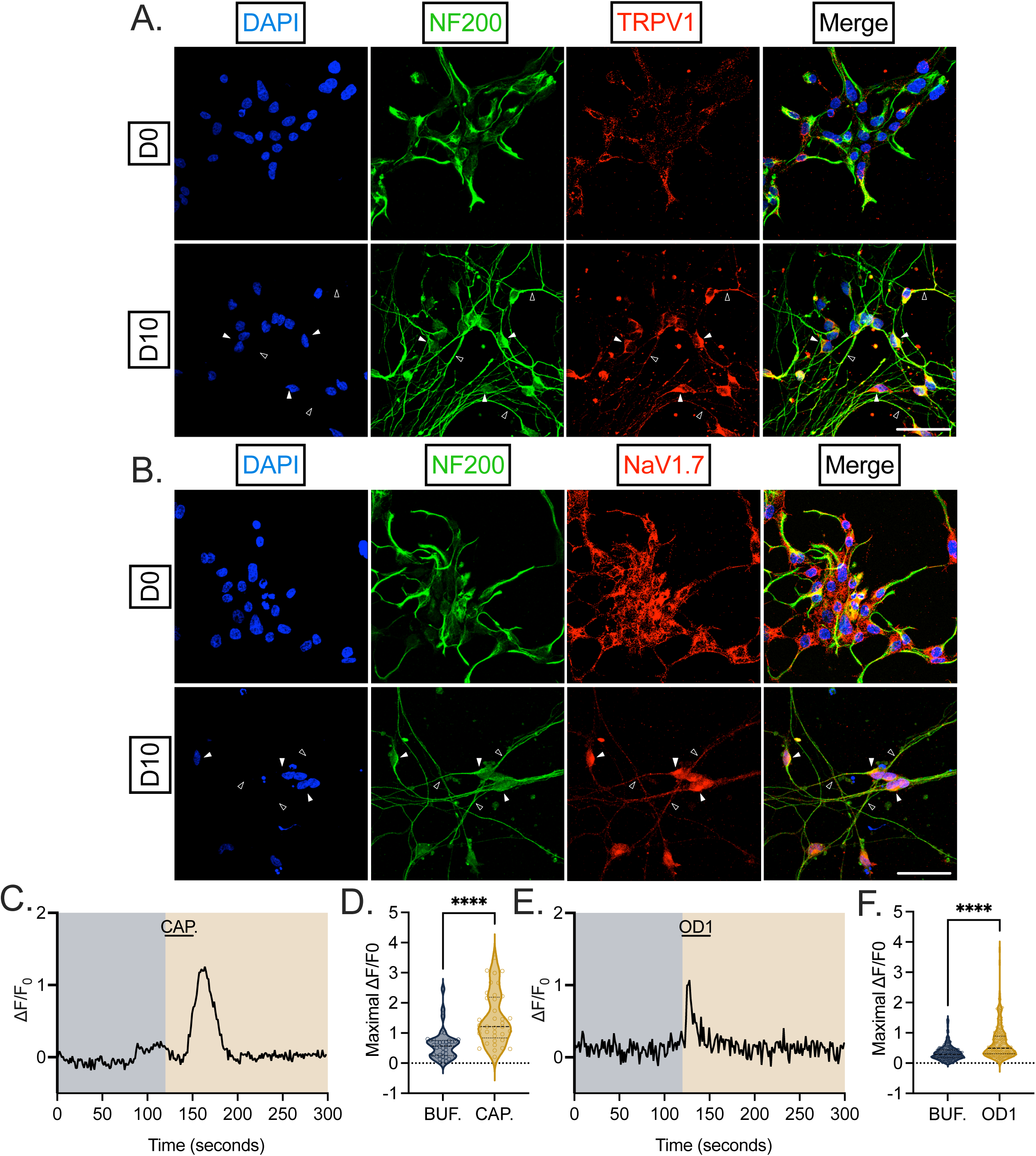
Key nociceptive ion channel abundance, localization, and activity in HD10.6 cells. **A**&**B**) D0 and D10 HD10.6 cells were fixed and stained for DAPI (blue), NF200 (green), and nociceptive ion channel TRPV1 (**A**) or NaV1.7 (**B**) in red. Representative images shown from three independent experiments. Scale bar 50 μm; filled arrows indicate examples of signal at soma whereas empty arrows indicate signal within axons. **C**-**F**) Calcium influx in response to buffer only (BUF., blue) as well as during and following 10 μM TRPV1 agonist capsaicin (CAP., **C**-**D**, orange) or 1 μM NaV1.7 agonist OD1 (**E**-**F**, orange) was recorded from differentiated HD10.6 cells with Ca^2+^ indicator Fluo-4AM (5 μM). Buffer was applied for 2 minutes and followed by the application of capsaicin or OD1 for 30 seconds. After up to 4 additional minutes of buffer, 50 mM KCl was applied for 5 seconds at the termination of the experiment to verify cell viability. Representative traces in response to respective stimuli shown (**C**&**E**), as well as maximal normalized intensity (**D**&**F**). Replicates from three independent experiments. The total number of cells analyzed per treatment was: 37 for CAP. and 213 for OD1. Statistical comparisons were made using Wilcoxon signed-rank test due to non-normality. *, p<0.05; **, p<0.01; ***, p<0.001; ****, p<0.0001.

There are nine isoforms of voltage-gated sodium channels (NaV1.1-1.9), and four isoforms expressed within the DRG (NaV1.3, NaV1.7-1.9) are implicated in the transmission of pain signals by epidemiological patient data and animal models^19^. Over the last few years, special attention has been paid to NaV1.7 for its potential therapeutic utility in small molecule or gene therapy form for a variety of chronic pain states^20–23^. Therefore, we investigated the expression and function of NaV1.7 within the HD10.6 cells and found that NaV1.7 was abundantly expressed along the soma and the axon in all differentiated HD10.6 cells (**Fig. 2B**). NaV1.7 protein has previously been observed in HD10.6 cells differentiated under a different maturation protocol^13^. We next probed the specific activity of NaV1.7 through calcium imaging with NaV1.7 agonist OD1. Scorpion toxin OD1 enhances NaV1.7 activity by inhibiting channel inactivation and increasing peak current^24,25^, and intraplanar injection of OD1 into mice elicits spontaneous pain behavior^26^. In response to OD1 exposure, we detected a transient and significant increase in Ca^2+^ influx in differentiated HD10.6 cells (**Fig. 2E-F**). Therefore, differentiated HD10.6 cells abundantly express characteristic nociceptive ion channels implicated in pain states, and these channels are responsive to direct stimulation.

### Electrophysiological properties of HD10.6 cells

We next performed whole-cell patch clamp electrophysiology, including current clamp and voltage clamp, on differentiated HD10.6 cells (**Figure 3**) to outline an electrophysiological profile of the cells and compare it to previously published parameters on human DRG neurons and nociceptors. C-fiber nociceptors are classified as “small diameter” DRG neurons^8,27^. Accordingly, the average diameter of the HD10.6 cell soma measures 19.3 ± 0.5 μm (**Fig. 3B**), which is slightly smaller than what is observed in post-mortem human DRG cultures^28^ but interestingly similar to that of nociceptor-like human induced pluripotent stem cells^29^. For D10 HD10.6 cells, the mean capacitance was 19.3 ± 0.9 pF (**Fig. 3C)**, a value slightly lower than that observed within human DRG cultures^28,29^ likely due to the smaller soma diameter. Under current clamp, the resting membrane potential of differentiated HD10.6 cells was -49.5 ± 1.2 mV (**Fig. 3D**). Human donor nociceptors demonstrate a similar resting membrane potential, ranging from -40 mV to -60 mV^28,29^. Importantly, resting membrane potential remained stable during the recording of untreated HD10.6 cells (**Fig. 3E**). Healthy DRGs isolated from human patients also demonstrate a stable resting membrane potential, whereas DRGs isolated from painful dermatomes display spontaneous ectopic activity^30,31^.

**Figure 3:**
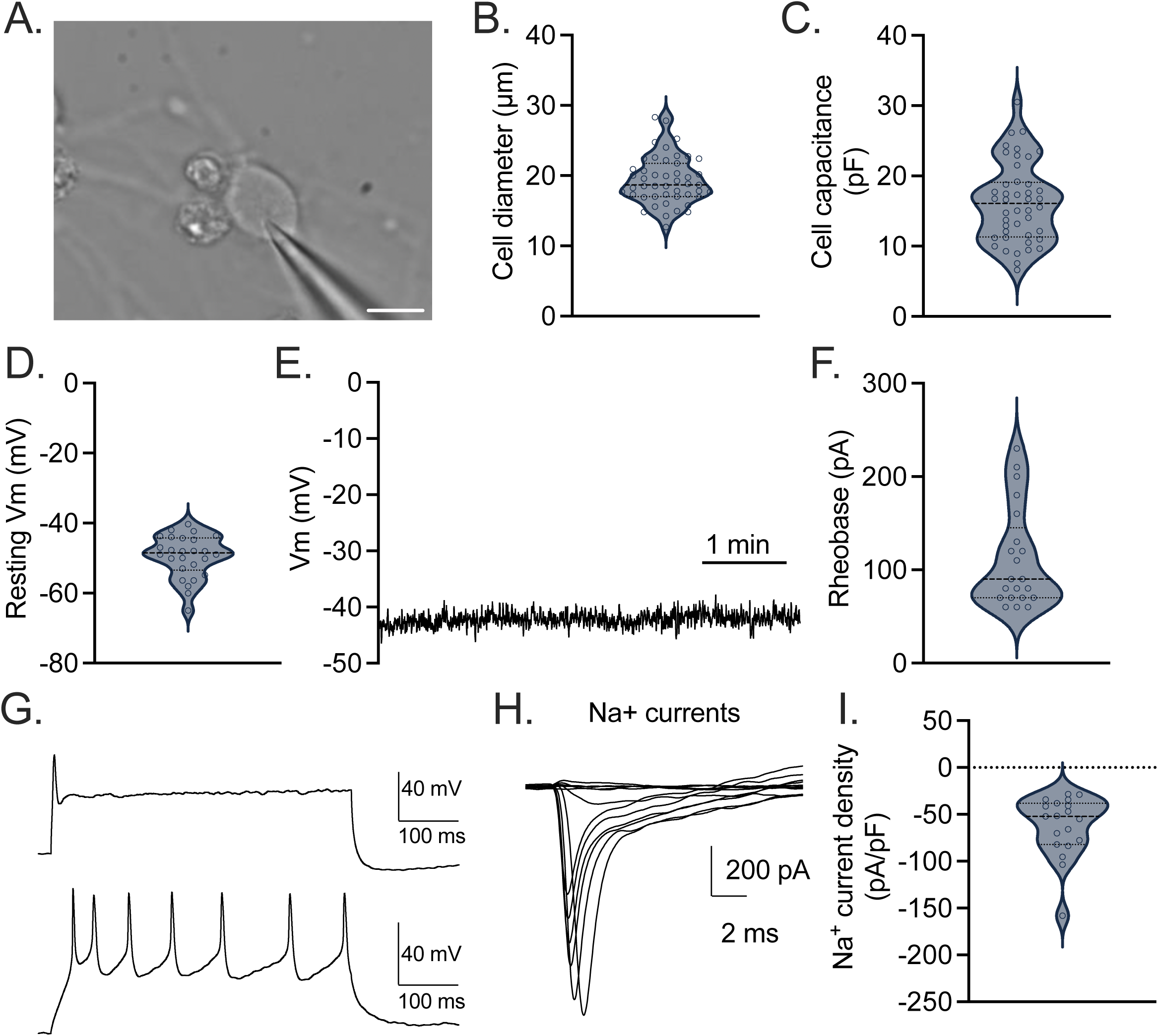
Electrophysiological parameters of differentiated HD10.6 cells. D10 HD10.6 cells under current clamp (**A-G**). **A**) An image of differentiated HD10.6 undergoing active whole-cell patch clamp. Scale bar 10 μm. **B**) Cell soma diameter. **C**) Cell capacitance. **D**) Resting membrane potential. **E**) Representative recording of stable resting membrane potential over time. **F**) Rheobase. **G**) Representative recording of single discharge and multiple discharge patterns following current injection. D10 HD10.6 cells under voltage clamp (**H**-**I**). **H**) Sodium currents elicited by depolarizations from a holding potential of −80 mV to test potentials from −70 mV to +40 mV with 10 mV increment. **I**) Na+ sodium current density. Replicates from three independent experiments.

The average rheobase was 111 ± 12 pA (**Fig. 3F**) and most differentiated HD10.6 cells investigated (92%) released a single discharge in current step recordings (**Fig. 3G**, top). However, a smaller proportion (8%) emitted two or more sequential discharges (**Fig. 3G**, bottom). Whether the cell emitted a single or multiple discharge(s) was independent of increasing injected current, reflecting an intrinsic commitment to the discharge pattern. In both human DRGs from donors or human nociceptor-like iPSCs, this commitment of both single and multi-firing cells is also observed^28,29^. Under voltage clamp, we identified sodium currents elicited by depolarizations, as previously observed^11^ (**Fig. 3H**), confirming their activity. The sodium current density was measured, averaging -62.5 pA/pF^32^ (**Fig. 3I**).

Notably, in previous works, HD10.6 cells were cultured with 25 μM forskolin (FSK)^13,33^. When we cultured HD10.6 cells with FSK in addition to the media described in the Methods, we found that the resting membrane potential was 30% higher (approximately -35 mV) than observed under incubation without FSK (data not shown). Under FSK incubation, action potentials were difficult to induce with current injections, suggesting the FSK incubation time (10 days) or concentration (25 μM) led to premature depolarization that led to the failure of action potential generation. In conclusion, differentiated HD10.6 cells, under our current described maturation protocol, have similar electrophysiological properties as nociceptive neurons from human patients.

### Afferent peptides and excitatory receptors expression and localization

The peripheral terminal of the nociceptor acts not only as a sensor of local (painful) stimuli, but also as an effector^34^. Nociceptors can store and release afferent peptides, such as CGRP and substance P which can have excitatory impacts on non-neuronal cells of the DRG through neurogenic inflammation and may directly feedback upon the nociceptor itself^35^. We therefore inquired whether HD10.6 cells expressed, at the protein level, characteristic afferent peptides (**Figure 4**). Indeed, we detected faint CGRP (**Fig. 4A**) and substance P (**Fig. 4B**) within undifferentiated HD10.6 cells, the signal of which grew more abundant following differentiation. The receptor for SP, neurokinin 1 receptor (NK1R)^36^, is expressed by vascular endothelial cells and immune cells, which contribute to the inflammatory aftermath of SP release. However, there is also evidence that nociceptors express NK1R on their surface^37^, suggesting an excitatory loop within the DRG wherein afferent peptides may feedback on the nociceptor may be at play. We inquired into the expression of NK1R within HD10.6 cells and found abundant NK1R staining in undifferentiated and differentiated HD10.6 cells (**Fig. 4C**). No signal was detected in control slides incubated without primary antibodies in any of these experiments. Therefore, differentiated HD10.6 cells abundantly express excitatory afferent peptides, as well as the Substance P receptor. Importantly, these data suggest that HD10.6 cells can be used to study the neurogenic contributions from and to human nociceptors.

**Figure 4:**
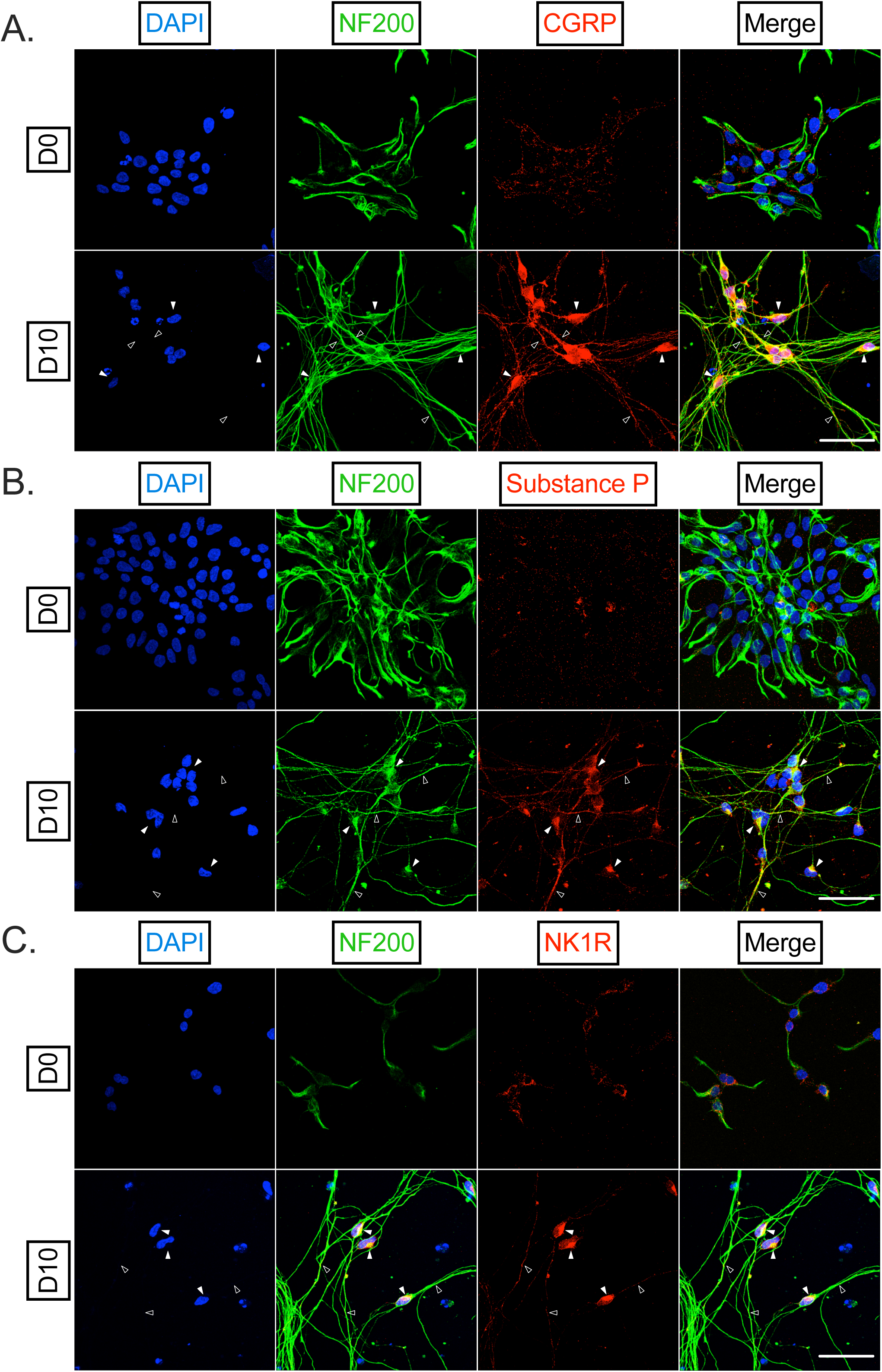
HD10.6 cells express key afferent peptides and related machinery associated with pain regulation. **A**-**B**) D0 and D10 HD10.6 cells were fixed and stained for DAPI (blue), NF200 (green), and afferent peptides CGRP (**A**) or Substance P (**B**), or Substance P receptor NK1R (**C**) in red. Representative images shown from three independent experiments. Scale bar 50 μm; filled arrows indicate examples of signal at soma whereas empty arrows indicate signal within axons.

### Functional opioid-induced analgesic signaling within HD10.6 cells

Opioid signaling has long been co-opted for significant pain relief in humans, although with substantial issues regarding dependency and addiction^38^. The role of opioid signaling within the peripheral terminal, in addition to the spinal cord, has become increasingly appreciated^39^. Rodent model neurons do not recapitulate the diversity and robustness of mu opioid receptors (MOR) within nociceptors^5^, nor do they entirely accurately recapitulate opioid use disorder^40^ at the transcriptional and epigenetic level, emphasizing the importance of human material in this study. We therefore inquired whether functional mu opioid signaling was present in differentiated HD10.6 cells (**Figure 5**). First, we stained both D0 and D10 HD10.6 cells for the mu opioid receptor (MOR). We found lowly detectable signal in D0 HD10.6 cells, but more abundant signal within the soma and axons of differentiated HD10.6 cells (**Fig. 5A**). To examine the functional capacity of opioid receptors in modulating electrical excitation, we tested whether mu opioids were able to quell evoked electrical excitement. Electrical excitation was initiated with 10 mM KCl and measured by calcium influx in the absence or presence of pre-treatment with the mu opioid selective agonist DAMGO. Pre-treatment with DAMGO for 2 minutes resulted in a strikingly attenuated calcium influx peak in response to KCl treatment (**Fig. 5B**), and the change in maximal calcium influx intensity was reduced by approximately 50% (**Fig. 5C**). This reduction in influx was not due to cell death, as treated cells responded to 50 mM KCl at the termination of the experiment. Importantly, the effect on influx by DAMGO was also reversed through the addition of opioid receptor antagonist naloxone, which slightly enhanced influx over control conditions as determined by maximal intensity (25% increase, **Fig. 5C**). The addition of DAMGO or the DAMGO/naloxone combination alone (in the absence of 10 mM KCl) did not induce calcium influx levels greater than those observed during perfusion with buffer. Therefore, differentiated HD10.6 cells display functional mu opioid signaling and can be used to model these phenomena.

**Figure 5:**
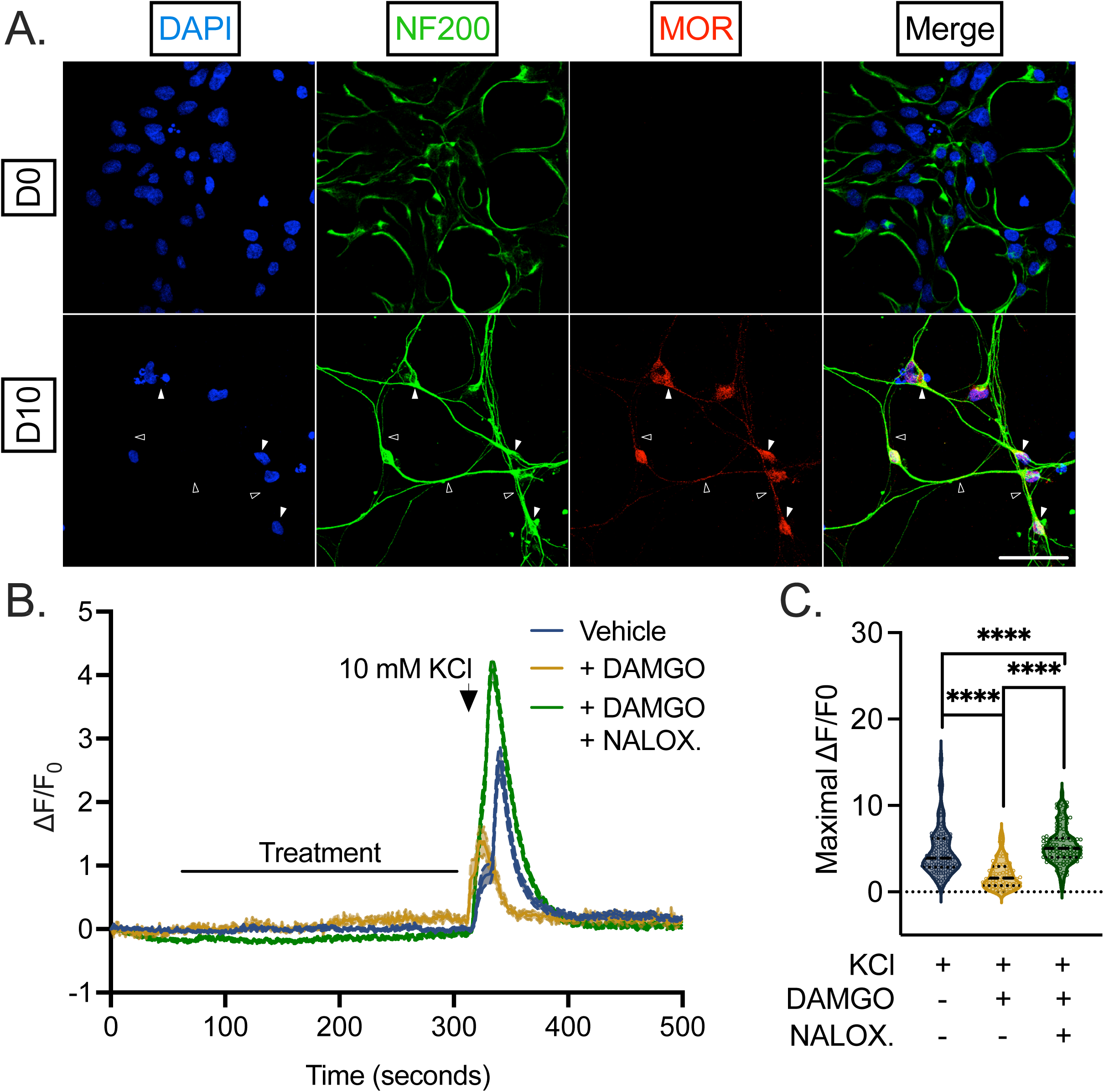
Functional opioid-induced analgesic signaling in human HD10.6 cells. **A**) D0 and D10 HD10.6 cells were fixed and stained for DAPI (blue), NF200 (green), and mu opioid receptor 1 (MOR1, red). Scale bar 50 μm; filled arrows indicate examples of signal at soma whereas empty arrows indicate signal within axons. **B**-**C**) Calcium influx in response to 10 mM KCl was recorded from differentiated HD10.6 cells with Fluo-4AM. Buffer was applied for 1 minute. Following an additional 4-minute perfusion of buffer with vehicle (blue), 1 μM DAMGO (orange), or both DAMGO and naloxone (NALOX.) at final concentrations of 1 μM and 10 μM, respectively (green), a 10-second application of 10 mM KCl was applied. After up to 4 additional minutes of buffer, 50 mM KCl was applied for 5 seconds at the termination of the experiment to verify cell viability. Combined traces (mean ± SEM, **B**), as well as maximal normalized intensity (**C**) in response to respective treatments. Replicates from three independent experiments. The total number of cells analyzed per treatment was 132 for vehicle, 72 for DAMGO, and 107 for DAMGO and NALOX. Statistical comparisons were made using an unpaired non-normal t-test (Mann-Whitney U test). *, p<0.05; **, p<0.01; ***, p<0.001; ****, p<0.0001.

### Peripheral sensitization: an inflammatory soup elicits and primes nociceptive signaling in HD10.6 cells

These data suggest that HD10.6 cells act as *de facto* human nociceptors and respond electrically to direct (anta)agonists of ion channels which mediate excitability. We next sought to perturb the excitability of the differentiated HD10.6 cells in a physiological manner that mirrors a painful condition to which a nociceptor might be subject (**Figure 6**). We therefore exposed HD10.6 cells to an “inflammatory soup” (IS)^41–43^, a mixture including bradykinin, prostaglandin E2 (PGE2), serotonin, adenosine triphosphate (ATP), and histamine, molecules released at the site of injury/inflammation as a result of the disruption of surrounding inflammatory cells. We aimed to model human “pain” using nociceptor excitability/activity as the best readout second only to behavior. First, we performed calcium influx live-imaging on the HD10.6 cells in response to transient (30s) IS application to assess whether this cocktail could directly enhance the excitability of the HD10.6 cells. In the scenario of acute pain, inflammatory mediators induce excitation and a pain state that subsequently resolves^44^. IS addition almost instantly induced a statistically significant and transient influx in calcium mirroring an action potential (**Fig. 6A-B**). Next, we queried whether transient IS application would prime the HD10.6 cells analogous to a process undergone by nociceptors known as peripheral sensitization^45^. This process, also known as hyperalgesic priming, is commonly used to study how acute pain transforms into chronic pain^46^. HD10.6 cells were pre-incubated with IS or vehicle control for a 4-hour period, following which the HD10.6 cells were thoroughly washed and prepared for calcium imaging where 10 mM KCl was used to initiate electrical excitation. HD10.6 cells pre-treated with IS responded to 10 mM KCl significantly more robustly than vehicle controls in terms of the fold change/potency of calcium influx within HD10.6 cells (**Fig. 6C-D**) as well as the number of HD10.6 cells that were responsive to the stimulus (**Fig. 6E**). Excitingly, these data suggest that the IS can be applied to HD10.6 cells to model pain in both the presence and following removal/resolution of a noxious stimulus. We verified this finding using the gold standard of physiology, whole-cell patch clamp. HD10.6 cells pre-incubated with IS for 4 hours had a resting Vm that was depolarized as compared to untreated cells (**Fig. 6F**, -IS: -49.5 ± 1.2 mV, n = 26; +IS: -45.3 ± 1.5 mV, n = 11; unpaired t-test, p < 0.05). The rheobase was also significantly lower for +IS conditions than that observed for untreated cells (**Fig. 6G-H**, -IS: 111 ± 12 pA, n = 21; +IS: 67 ± 7 pA, n = 11; unpaired t-test, < 0.05), and 30% of IS-treated cells showed multiple discharges in current step recordings. These data indicate that HD10.6 cells incubated with IS are more likely to fire action potentials than untreated cells. No difference in cell capacitance and cell soma diameter between the IS-treated and untreated cells was observed, which suggests that IS incubation did not induce significant cell swelling indicative of poor health (**Fig. 6I-J**). A significant difference in sodium current density was also demonstrated between the control and experimental groups (**Fig. 6K**). Importantly, we conclude HD10.6 cells undergo peripheral sensitization and can therefore be used to model both acute and persistent pain states.

**Figure 6:**
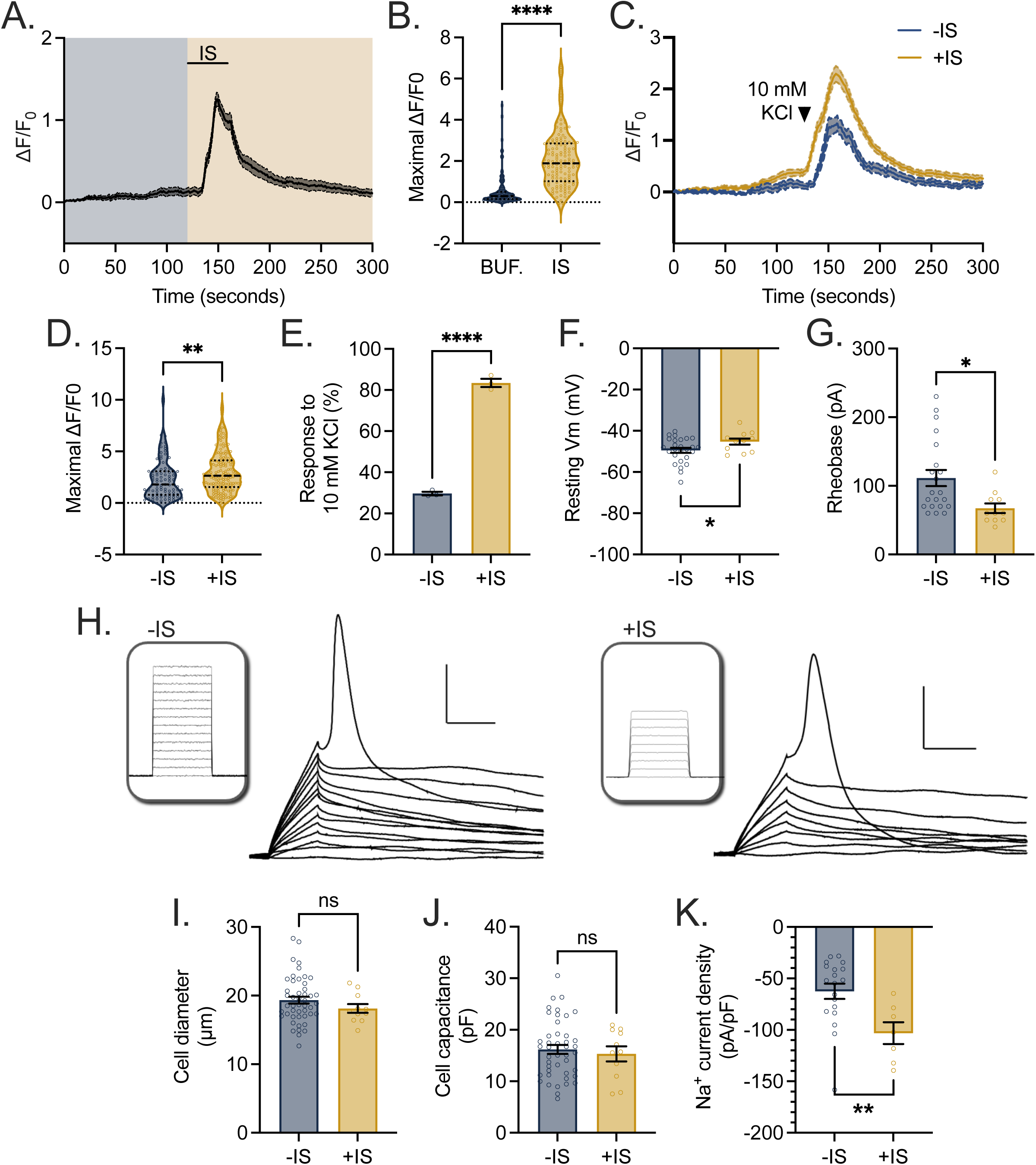
HD10.6 cells undergo peripheral sensitization in response to an inflammatory cocktail. **A**-**B**) Calcium influx in response to buffer only (BUF., blue) as well as during and following application of inflammatory soup (orange) was recorded from differentiated HD10.6 cells with Fluo-4AM. Buffer was applied for 2 minutes, and the inflammatory soup (IS) was applied for 30 seconds. After up to 4 additional minutes of buffer, 50 mM KCl was applied for 5 seconds at the termination of the experiment to verify cell viability. Combined traces (mean ± SEM) in response to stimuli shown (**A**), as well as maximal normalized intensity (**B**). Replicates from two independent experiments. The total number of cells analyzed was 133. **C-K**) D10 HD10.6 cells were incubated with vehicle (-IS) or IS for 4 hours. (**C**-**E**) Following incubation and thorough washing, calcium influx in response to buffer and a 10-second application of 10 mM KCl was recorded. After up to 4 additional minutes of buffer, 50 mM KCl was applied for 5 seconds at the termination of the experiment to verify cell viability. Combined traces (mean ± SEM) in response to treatment and stimuli shown (**C**), as well as maximal normalized intensity (**D**) and the percentage of HD10.6 cells responsive to 10 mM KCl (**E**). Replicates from two independent experiments. The total number of cells analyzed per treatment was: 87 for -IS and 169 for +IS. (**F**-**J**) Following incubation and thorough washing, whole-cell patch clamp was performed on HD10.6 cells. **F**) Resting membrane potential. **G**) Rheobase. **H**) Representative rheobase recordings. Insets: I-command, 10 pA increment, 10 ms duration. Scale bar: 20 mV, 10 ms. **I**) Cell soma diameter**. J**). Cell capacitance. (**K**) Following incubation and thorough washing, voltage clamp recordings were performed on HD10.6 cells. Na+ sodium current density is shown. Statistical comparisons were made using an unpaired t-test or Wilcoxon signed-rank test dependent on normality of the data. *, p<0.05; **, p<0.01; ***, p<0.001; ****, p<0.0001.

### The peripheral terminals of HD10.6 cells show AAV uptake, transport, and soma transfection

One of the limitations of many current *in vitro* nociceptive models is the disorganization of these cultured primary or immortalized DRG neurons within wells. Regarding the primary afferent, the nociceptor soma resides in the DRG and the peripheral axon terminals which detect and respond to painful stimuli innervate the periphery a distance away. These are distinct physiological environments in that the soma and peripherals axons are in contact with differential local/circulating inflammatory products and non-neuronal cells. The intrinsic biology of the soma and axon of any neuron is also distinct in terms of signaling integration, action potential initiation, and the distribution and influence of ion channels^47^. We therefore optimized the culture of HD10.6 cells into microfluidic chambers in which we can fluidically isolate, stimulate, and study the peripheral terminal and soma to effectively mirror the anatomy of a human patient (**Figure 7**).

**Figure 7:**
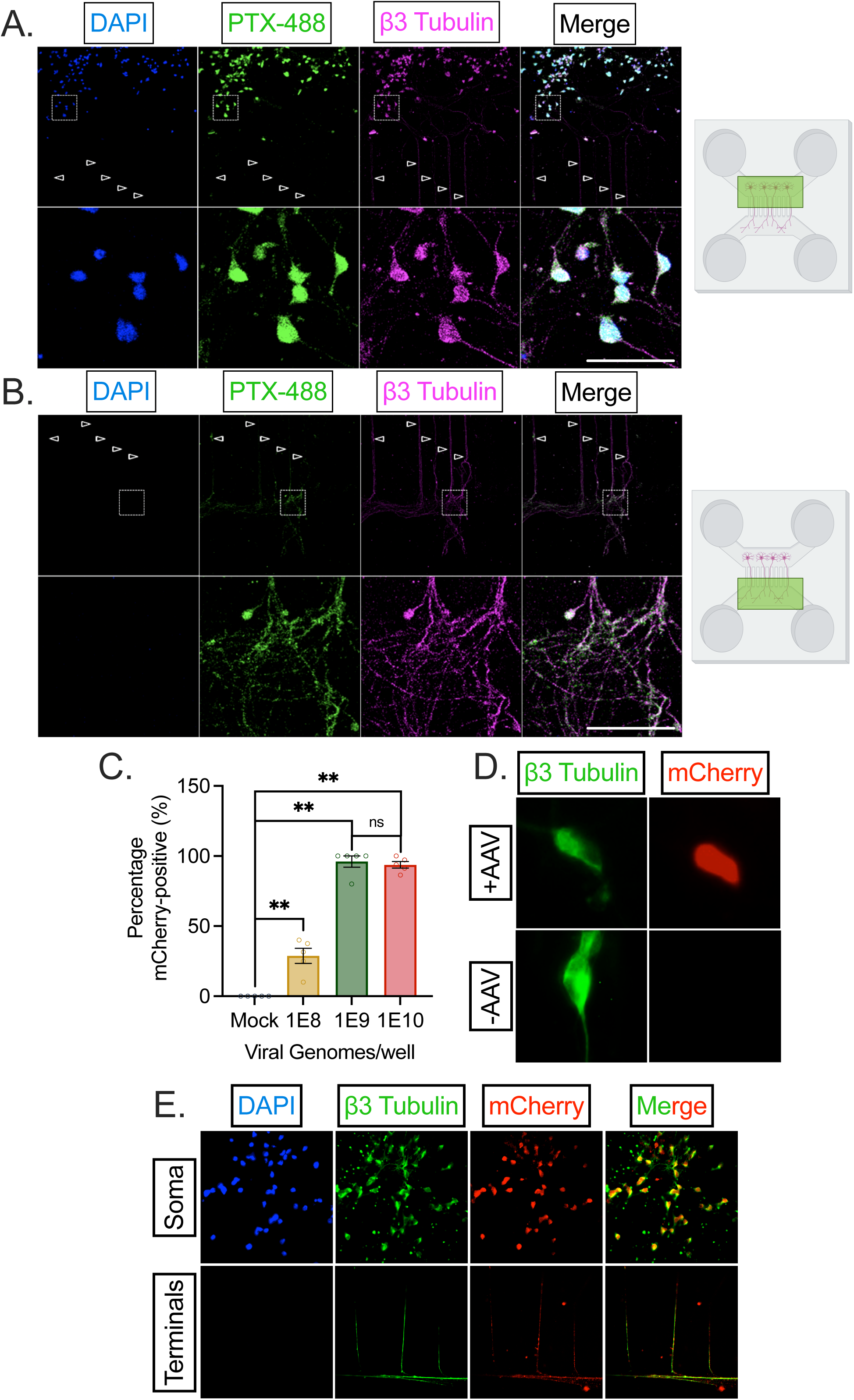
Culture of HD10.6 cells into microfluidic devices to study peripheral administration. **A**-**B**) D0 HD10.6 cells were seeded into the soma compartment of a microfluidic chamber, and media was changed to differentiation media 24 hours post-seeding. A 2X nerve growth factor (NGF) gradient was used to guide axons towards the axon-only compartment. At D14, the cultures within the chamber were fixed and stained for neuronal marker β3 tubulin (magenta), marker for NaV1.7 PTX-488 (green), and DAPI (blue). Empty arrows point to axons traveling through microgrooves. **A**) captures a representative image of the soma compartment and axons within the proximal end of the microgrooves, whereas **B**) shows the axon-only compartment and distal microgrooves. Scale bar 50 μm. Respective compartments are highlighted in green within the right-hand animated microfluidic chambers; images of microfluidic chambers created with BioRender.com. **C**-**D**) D10 HD10.6 cells in cell culture wells were infected with AAV9-mCherry. 3 days following infection, cells were fixed and stained for β3 tubulin (green), mCherry (red), and DAPI (blue) The percentage of mCherry+ HD10.6 cells based on the total number of cells identified by DAPI and β3 tubulin-positive staining is demonstrated in **C**. Counts were performed under blinded conditions. A representative image of AAV9-mCherry-treated (1E9, top) and untreated (bottom) HD10.6 cells at this time-point is shown in **D**. **E**) 3 days following AAV9-mCherry infection in the peripheral compartment of HD10.6 cells cultured in microfluidic chambers, the cultures were fixed and stained for β3 tubulin (green), mCherry (red), and DAPI (blue). Representative image of soma (top) and peripheral terminal compartments (bottom) demonstrate mCherry signal within the soma and axons of HD10.6 cells. Replicates from three independent experiments; statistical comparisons were made using an unpaired non-normal t-test. *, p<0.05; **, p<0.01; ***, p<0.001; ****, p<0.0001.

Within the microfluidic device, there are separate soma and axonal compartments which are connected by 2 μm microgrooves. These microgrooves are large enough for the passage of neuronal axons but not cell bodies. D0 HD10.6 cells were seeded into the soma compartment of the microfluidic chamber and differentiated 24 hours later. A nerve growth factor (NGF) gradient was used to guide axons from the soma compartment through microfluidic grooves towards the axon-only compartment. By 5-8 days post-differentiation, axonal extensions sprouted into the axon-only compartment. The length of the microgrooves in this system is 450 μm, therefore reflecting a tremendous capability of HD10.6 cells to sprout. At D15, the cultures within the chamber were fixed and stained for neuronal marker β3-tubulin, marker for NaV1.7 ATTO488-ProTx-II (PTX-488)^48^, and DAPI. Within the soma compartment, we demonstrate the confinement of viable cell bodies through the presence of rounded soma staining positive for both PTX-488 and DAPI (**Fig. 7A**). Robust dendritic branching is apparent through the microgrooves (empty arrowheads) and in the axon-only compartment representing the peripheral terminal (**Fig. 7B**).

This anatomical distinction is advantageous to study not only the physiology of pain states, but also the therapy. The use of gene therapies, such as adeno-associated virus (AAV) and antisense oligonucleotides (ASO)^49^, to treat chronic pain states is a rapidly advancing field. While these vectors are commonly delivered through an intrathecal injection (analogous to treatment in the soma compartment of the HD10.6 cells within the microfluidic chamber), their peripheral administration is being increasingly investigated and applied^50,51^ due to efficacy, convenience, and specificity. Peripheral administration of the therapy, such as through subcutaneous injection, occurs at the periphery from where the AAV/ASO must infect the peripheral terminal endings and undergo retrograde transport to travel to the soma of the nociceptor. We recognized the suitability of the system of HD10.6 cells cultured within the microfluidic chambers to examine whether infection within the peripheral terminal would be a plausible route of inoculation in human material. Following the optimization of viral titer for mCherry-AAV9 uptake (**Fig. 7C-D**), the peripheral terminal was infected with 1E9 vg^52^. Three days later, chambers were fixed and stained for mCherry. mCherry staining, detected in both the soma and axons, was present in 100% of HD10.6 cells as identified by the presence of β3-tubulin^11,12^ (**Fig. 7E**). Therefore, AAV9 infection, retrograde transport, and expression proceeds within HD10.6 cells cultured in microfluidic chambers, having wide-ranging implications for future studies on not just mechanisms of pain signaling and chronification, but also its therapy.

## Discussion

Since the introductory publication of HD10.6 cells, it has not been concluded whether these neuronal cells are in fact nociceptors versus alternative DRG neuron subtypes^11^, and few other groups have characterized or employed these cells^12,13,33,53^. We systematically characterized the presence of nociceptive machinery within the neuronal soma and axons at the protein level, including TRPV1, NaV1.7, CGRP, Substance P, and NK1R. We verified through calcium influx assays that differentiated HD10.6 cells respond to direct agonists of TRPV1 and NaV1.7 and characterized the electrophysiological profile of differentiated HD10.6 cells through whole-cell patch clamp. We found that mu opioid administration robustly diminishes electrical activity, and that this phenomenon can be recovered with the addition of naloxone, demonstrating functional mu opioid signaling as observed in human patients. Importantly, we also found that HD10.6 cell excitability can be enhanced through the addition of an inflammatory soup, leading to both the direct activation of excitable events, as well as the peripheral sensitization process that is essential to developing persistent pain states. Finally, we optimized the culture of HD10.6 cells into microfluidic chambers to enable the physiological investigation of the soma versus the peripheral terminal. We found that HD10.6 cells successfully take up AAV through infection of the peripheral terminal, mirroring peripheral therapeutic administration. Altogether, we have developed the use of HD10.6 cells into a model of human nociception in a dish that can be employed for the investigation of molecular mechanisms of pain signaling and its therapy.

This model of nociceptive signaling in HD10.6 cells carries both inherent advantages and limitations. While we employ whole-cell patch clamp and calcium imaging within the HD10.6 cells to study excitability as a readout of the HD10.6 cell pain response, the ultimate readout of pain is behavior. As behavior can only be measured *in vivo*, we do not assert that DRG neuron work within animal models such as rodents should be discontinued; these *in vitro/in vivo*, human/animal model systems should be considered in parallel with their respective strengths. Further, this is not the first-time human material has been used to model or study nociception and pain. The Gareau, Dougherty, and Price labs, among others, have emphasized isolating and experimenting with human DRGs from patients, taken secondary to their removal by surgical necessity^54^. However, given their source, these DRG neurons are difficult to acquire with regularity and are likely diseased, necessitating their removal. Both a virtue and limitation of these human harvested tissues in DRG cultures is their inherent diversity based on epidemiological patient parameters like sex, age, and the potential of previous injury or infection. Human nociceptor-like neurons have also been created through induced pluripotent stem cell (iPSCs) protocols^55–57^ . The use of iPSCs is advantageous in that it can be used to create neurons with a customized genetic makeup from a patient. However, the fate of stem cells is largely promiscuous or prone to change with any minor adjustment to maturation media composition^58^. HD10.6 cells, on the other hand, exhibit fate restriction and retain their neuronal phenotype even when cultured in medias that support the differentiation of cells into Schwann cells or smooth muscle cells^11^. The reproducible nature, proliferative ability, and quick differentiation time of HD10.6 cells also make this cell line suitable for high throughput screening. Their embryonic nature also diminishes the probability of complications and alterations from external environmental stimuli (infection, injury). Concurring, we detect little ectopic activity in differentiated, but untreated HD10.6 cells (**Fig. 3E**).

Recently, an elegant human DRG organoid system was constructed following the single-cell spatiotemporal analysis of human embryonic DRGs at the transcriptomic level^59^. This system recapitulates all cell types present in the DRG, including multiple neuronal subtypes, glia, and Schwann cells, but it also requires regular access to human embryonic DRGs as well as a 90 day-differentiation period, leading to concerns over feasibility. The HD10.6 cell system does not carry all the components of a DRG, which is both a deficiency and benefit in that it provides a reductionist approach to studying the nociceptor. Nociceptor-intrinsic signaling can be analyzed without complication, and the impact of individual additional cell types or immune components can be deciphered. In our studies, we use a physiological inflammatory cocktail containing products released under inflammation or injury that excite and sensitize the nociceptor. Under stimulus, PGE2 is released from immune cells, epithelial cells, and stromal cells; ATP from macrophages, neutrophils, and endothelial cells; and serotonin/histamine/bradykinin from mast cells and basophils. In this way, we are accounting for the contributions of non-nociceptor cells to pain sensation and signaling, and we can further dissect the individual contributions of each soup “ingredient” to altering HD10.6 excitability in future studies.

Our finding that HD10.6 cells can respond to this soup is perhaps one of the most impactful findings of our study in that it shows this system can not only be used to study the nociceptor immediately following direct addition of painful stimuli, but that it can mirror the peripheral sensitization process so intricately tied to the transition from acute to chronic pain states. This speaks to the potential utility of these cells to model differential chronic pain states. In our studies, we noted excellent cell viability following 30+ days post-maturation (data not shown). To enhance the translatability of this system in general and to the study of chronic pain states, we cultured HD10.6 cells into a microfluidic chamber that mirrors the anatomy of the peripheral terminal. Our modification enables the opportunity to co-culture different non-neuronal cells within the periphery (peripheral terminals) or DRG (soma compartment) and study their interactions with nociceptors. It also enables differential treatment with viruses or factors released from these other cells. The anatomically distinct system can also be used to mirror delivery methods for pain therapeutics. Intrathecal deliveries of drugs or genetic vectors can be modeled through the administration of the therapy into the soma compartment. Locally targeted therapies, including topical ointments or genetic vectors through subcutaneous injection, can be administrated into the pseudo-human periphery through incubation in the peripheral terminal axon end. In this work, we found that AAV9-mCherry is successfully taken up in HD10.6 cells following infection of the peripheral terminal. Limited work has been performed following subcutaneous injection administration of AAV therapies^50,51^ despite its advantages^60,61^. We put forward this model as a general human screening system for the efficacy of both soma and axon/terminal administered therapeutics. We hope to further adapt this system to pathological pain states to more accurately mirror the peripheral terminal of a human patient.

## Acknowledgments

We thank Dr. Chengbiao Wu and Dr. William Mobley for their advice on microfluidic chamber use. We thank Dr. Brian Brenner and Dr. Lynn Kohan for their clinical feedback on this manuscript.

